# Evolution of maternal and early zygotic transcript regulation across Drosophila

**DOI:** 10.1101/2021.10.28.466359

**Authors:** Charles S. Omura, Susan E. Lott

## Abstract

The complements of mRNAs in early embryonic development are crucial for setting up developmental trajectories across all animals. The earliest stages of development are regulated by mRNAs deposited into the egg by the mother, until the zygote can become competent to transcribe its own genome. Previously, we showed that the set of maternally deposited and early transcribed zygotic mRNAs in *Drosophila* are generally conserved across species, but with some notable variation. We also showed that a majority of regulators of these two types of transcripts are shared. In this study, we examine the differences in regulatory motifs associated with maternal deposition and early zygotic transcription across species of *Drosophila*. For maternal transcripts, while the regulators are mostly conserved, we find the *Drosophila pseudoobscura* species subgroup appears to contain numerous novel regulatory motifs unique to these species. These novel motifs are enriched in transposable elements exclusive to this group. As this species group had been previously identified as having an exceptional amount of divergence in early embryonic transcripts, this change in regulation may be responsible. However, transcripts that are present at the maternal stage only in these species are equally enriched in novel (group-specific) and conserved binding sites, so the novel regulation is not the sole cause of regulatory divergence in these species. At the zygotic stage, we observe a wide variety of species-specific motifs. Additionally, at both stages we observe motifs conserved across species having different effects on gene expression in different species, and regulating different sets of genes in different species. By examining changes in motif content across species, we find that changes in motif content alone is generally insufficient to drive gene expression changes across species.

## Introduction

The beginning of animal development is an especially dynamic period, where an egg with a single diploid nucleus after fertilization starts the process of building an entire individual. These early stages are exceptionally important, as these processes set up trajectories for the rest of development. The processes involved, such as rapid cleavage cycles and axial determination are largely conserved across animals. One such critical feature of early development, conserved across metazoa, is that early development is carried out by two different genomes: that of the mother and that of the zygote (Vastenhouw, Cao, and Lipshitz 2019; Tadros and Lipshitz 2009; Baroux et al. 2008). The very earliest stages of development are controlled by maternally deposited mRNAs and proteins, until the zygotic genome becomes competent to transcribe its own genome. This involves a highly regulated and coordinated handoff of developmental control between the maternal and zygotic genomes, known as the maternal to zygotic transition (MZT).

The transcripts provided to the embryo by the mother and transcribed by the zygote are also produced in strikingly different transcriptional contexts. In *Drosophila,* the maternal transcripts are produced by polyploid support cells called nurse cells during oogenesis; nurse cells rapidly produce large quantities of RNA (>100ng total RNA deposited per oocyte) representing a large proportion (up to 75%) of the genome (Vastenhouw, Cao, and Lipshitz 2019). Maternal transcripts are also subject to considerable post-transcriptional regulation, such as actively regulated stabilization (Cui et al. 2013; Eichhorn et al. 2016), degradation (Tadros et al. 2007; Bashirullah, Cooperstock, and Lipshitz 2001), and localization within the embryo (Medioni, Mowry, and Besse 2012; Vazquez-Pianzola et al. 2017). Presumably the role of post-transcriptional regulation is especially significant for maternal transcripts as regulation at the transcriptional level is not possible during the stage of embryogenesis when only maternal transcripts are present. In contrast, transcription of zygotic genes can require a considerable degree of precision, as some expression patterns of these genes are highly spatially and temporally restricted. For example, the well-studied segmentation patterning gene *even-skipped*, which has been a model for seminal work on how gene expression works in a complex eukaryotic system, is expressed at a particular stage of development in a seven stripe pattern along the anterior-posterior axis (Arnosti et al. 1996; Small et al. 1991; Levine 2010). While produced in considerably different transcriptional environments, the transcripts provided to the embryo by the mother and those later transcribed by the zygotic genome work together to carry out highly conserved processes of early development.

While many of the processes of early development are conserved across animals, the extent to which the gene products involved are also conserved is less well understood. For the earliest stages before and after zygotic genome activation, different complements of RNA transcripts are present across well-studied model species such as mouse, zebrafish, and *Drosophila* (Heyn et al. 2014). Considering the vast divergence time between these species, it is unclear from these comparisons how differences in early transcriptomes evolve. Previous work (Atallah and Lott 2018; Chenevert 2022) pursued this by examining transcriptomes of early embryos across 14 *Drosophila* species, and found high degrees of conservation in early zygotic and especially in maternally deposited mRNAs, with some notable exceptions.

In this study, we address the regulatory basis of evolved differences in maternal and zygotic transcriptomes across *Drosophila.* Given the importance of these early embryonic genes, exploring the regulatory mechanisms behind transcription in different species gives us insight as to how such traits evolve. We previously discovered a number of regulatory elements associated with maternal deposition and early zygotic transcription, where we also observed an overarching pattern of conservation, but with conspicuous deviations (Omura and Lott 2020). In this study, we examine how changes in regulatory binding sites relate to changing expression patterns for these critical stages. We found some lineage-specific motifs, with striking examples of differences concentrated in particular lineages at the maternal stage. We also found many well-conserved motifs, while they represent the same binding site, nonetheless seem to have differing effects on transcription across *Drosophila*. Given what we have discovered about motif content, we are still unable to broadly predict gene expression across species, and thus there are still additional factors yet to be determined.

## Results

In this study, we had two parallel goals: 1) to examine whether gene regulation had itself evolved in the maternal and early zygotic genomes that produce early embryo transcriptomes, and 2) to investigate the regulatory basis of evolved changes in gene expression of these same genomes. One primary method to address gene regulation was to discover motifs associated with maternal or early zygotic transcription. We applied this method to additional data from an RNA-Seq data set generated by a previous study (Atallah and Lott 2018), which provided us with transcript abundances across 14 species for two timepoints. One timepoint is at a stage of embryogenesis where all transcripts were derived from maternal deposition (stage 2, Bownes’ stages; (Bownes 1975; Campos-Ortega and Hartenstein 1985), the other after zygotic genome activation (late stage 5 or end of blastoderm stage). 11 of these species had genome annotations sufficient for this detailed motif analysis. We identified sequence motifs proximal to the transcription start site that had higher abundances in the maternally deposited or zygotically transcribed genes. We were then able to explore the occurrence of these motifs and associate them with transcript abundance in multiple species and found many examples where changes in motif content were associated with expression changes (Figure 1). Additionally, we evaluated interactions between motifs and a number of other factors commonly associated with gene expression, including gene length, distance to nearby genes, and strandedness.

**Fig 1:**
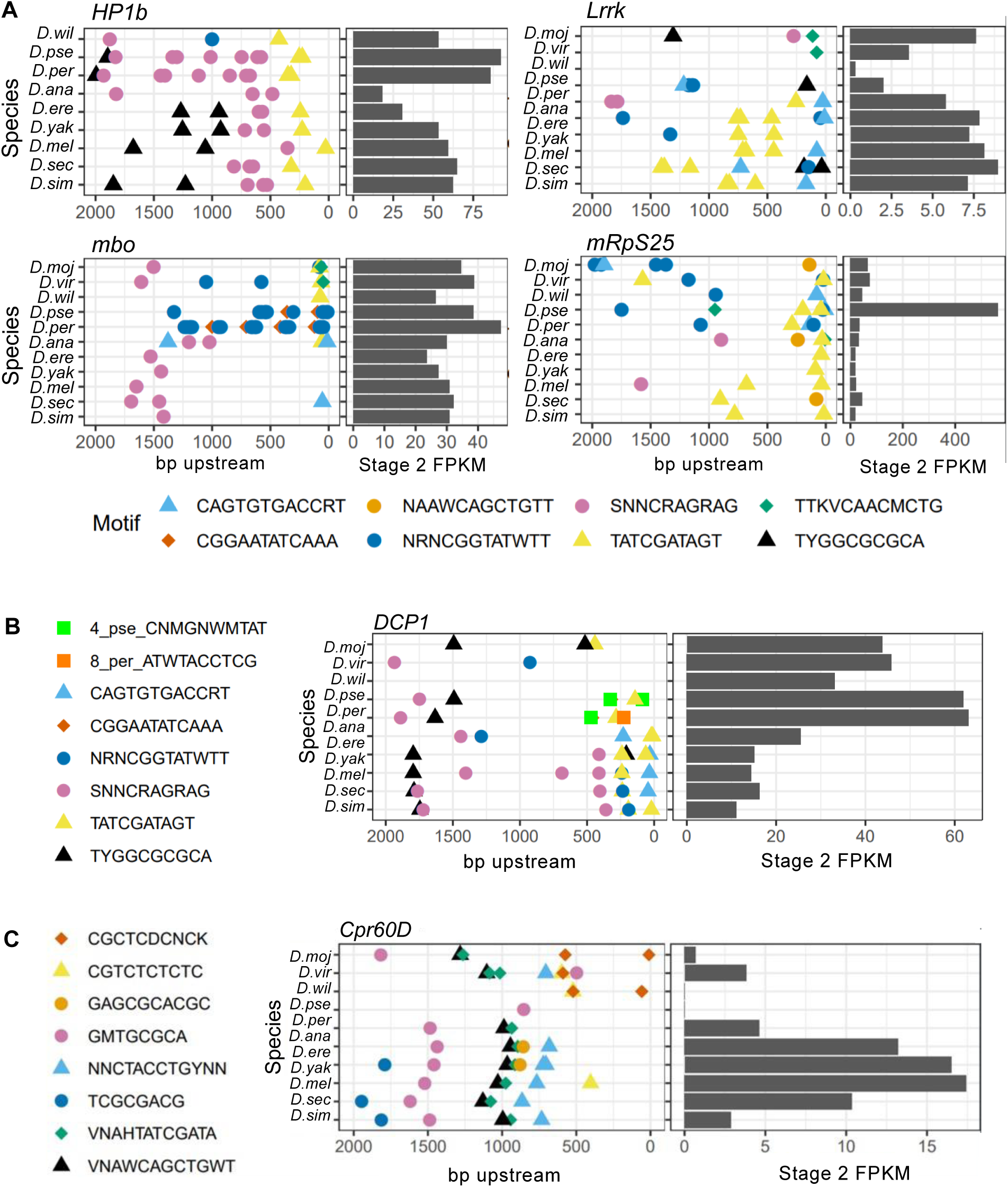
Changes in motif content are related to changes in expression. For both maternal and zygotic genes, we found numerous examples of genes whose transcript abundance and motif content have both evolved. Each left panel represents the 2kb upstream region of the gene, with the TSS at 0. Each colored symbol represents one motif. Each right panel indicates the measured transcript level (in FPKM) for each species. (A) Examples showing changes in conserved motifs at stage 2 that are associated with different transcript levels. (B) An example of a gene with changing transcript level associated with species-specific motifs in the upstream region at stage 2. (C) An example of changing transcript level associated with changing motifs at stage 5. Species are *D. melanogaster (Dmel), D. simulans (Dsim), D. sechellia (Dsec), D. yakuba (Dyak), D. erecta (Dere), D. ananassae (Dana), D. persimilis (Dper), D. pseudoobscura (Dpse), D. wilistoni (Dwil), D. virilis (Dvir), and D. mojavensis (Dmoja)*.

### Evidence for lineage-specific regulation

One fundamental question as to how gene expression is regulated in early development and how it evolves is to what extent the regulators are shared across species. Previously we demonstrated that there are a large number of regulators that are shared across species (Omura and Lott 2020). Here, we ask whether we can find evidence for lineage-specific regulators, that is, those specific to particular lineages or individual species. For this purpose, we surveyed the genomes of 11 *Drosophila* species for motifs (see Methods). In each species, we used HOMER (Heinz et al. 2010) to find a set of motifs that are enriched in the upstream region of either maternally deposited genes (stage 2) or of genes that are expressed after zygotic genome activation but not maternally deposited (zygotic-only genes, see Methods for definition; a subset of transcripts present at late stage 5). We performed this analysis separately in each species to identify motifs from all species.

With the motifs identified from all species in hand, we asked which motifs were acting in a lineage-specific manner. For each species, we constructed a linear model to relate the motif abundance with gene expression. We then compared the magnitude of the effects of each motif across species. We ranked motifs on how important the species term was on the model (see Methods). Many of the high-ranking motifs in maternal genes (from stage 2 data), exemplified by those represented in Figure 2A, showed high species-specific effects, with most motifs having a strong effect in only in the pseudoobscura subgroup species in the study, *D. persimilis* and *D. pseudoobscura*. Of the top 30 motifs, 11 were associated with increased transcript abundance only in *D. persimilis* and *D. pseudoobscura,* while another 15 displayed the strongest effect in *D. persimilis* and *D. pseudoobscura* and a weaker association with transcript abundance in other species. Motifs with a strong *D. persimilis* and *D. pseudoobscura* -specific association with transcript abundance occurred most frequently in these two sister species (Figure 2B).The differences between this subgroup and the reset of *Drosophila* is striking, given that these species are only diverged from *D. melanogaster ∼*30 million years ago (MYA), yet *D. melanogaster* and *D. virilis,* for example, have a longer divergence time (∼50 MYA) (C. A. M. Russo et al. 2013; C. A. Russo, Takezaki, and Nei 1995; Obbard et al. 2012) but show a much more similar pattern in motif content. While this pattern of *D. persimilis* and *D. pseudoobscura* sharing maternal motifs that differ from the rest of the species was the most common among motifs displaying species-specific effects, we also found examples of motifs that had different effects between *D. persimilis* and *D. pseudoobscura* (Figure 2C,D). After zygotic genome activation, we found no such bias of motifs associated with any particular species or lineage (Figure 2E,F,G,H). Instead we observed only sporadic examples of lineage-specific motifs in different individual species or groups.

**Fig 2:**
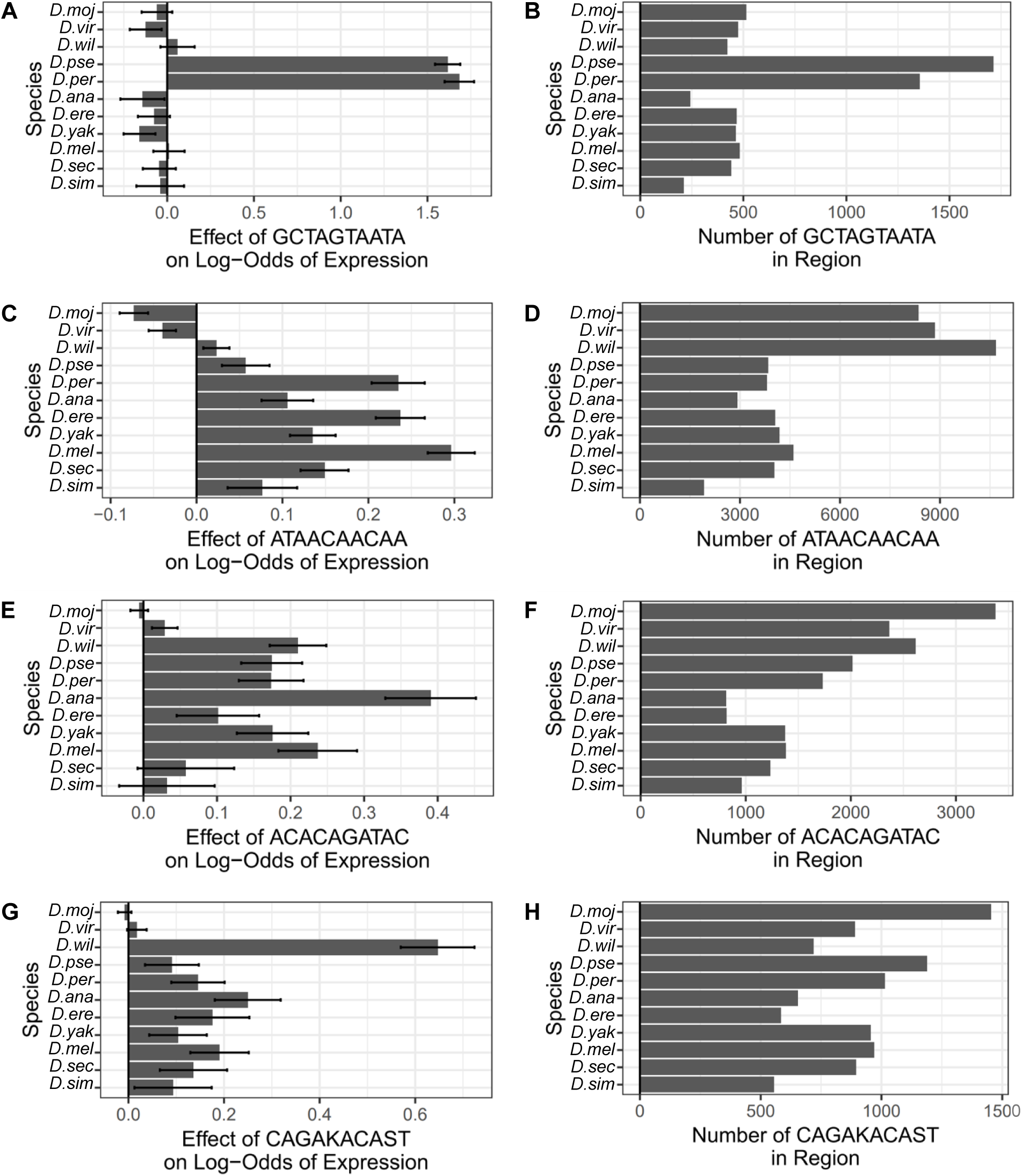
Discovered motifs can act in a species-specific manner. We identified a number of motifs that have species-specific effects. (A) A representative of a maternal gene motif that has a strong association with transcript abundance of target genes in *D.persimilis* and *D.pseudoobscura*, but not other species. (B) The same motif featured in (A) is present in all species, but is more frequent in the putative regulatory sequence of *D.persimilis* and *D.pseudoobscura*. (C) An example of a maternal stage motif that displays different effects even between closely related species, including *D.persimilis* and *D.pseudoobscura*. Interestingly, this motif apparently is associated with decreased transcript level in *D. mojavensis* and *D. virilis.* (D) This motif, seen in (C) is more common in those species where it has the smallest effect on transcript abundance. (E) An example of a zygotic stage motif with different effects on different species, with a particularly large effect on the odds of transcription in *D. ananassae*. (F) This motif (also in E) is least common in *D.ananassae.* (G) An example of a zygotic gene associated motif with different effects in different species, with a particularly large effect in *D.willistoni*. (H) Motif frequency does not appear to be correlated with putative motif effect for this motif.

### Species-specific motifs are not specifically associated with species-specific transcripts

Combined with our previous data (Omura and Lott 2020), we had identified a set of motifs that is species-specific and a set that is conserved. We asked whether these two sets of motifs occur within regulatory regions of separate sets of genes or whether they overlap within regulatory regions of the same set of genes. To this end, we measured the correlation of motif counts within regulatory regions of genes between all pairwise combinations of motif types at stage 2 (Figure 3A). We found that while there is a slightly higher correlation in location among the species-specific motifs than there is among the conserved motifs, the positive correlation between species-specific and conserved motifs suggests that they also occur within regulatory sequences of the same genes. This indicates that the species-specific motifs tend to co-occur in regulatory regions of the same genes with each other but are also distributed throughout the genome in a manner similar to the conserved motifs.

**Fig 3:**
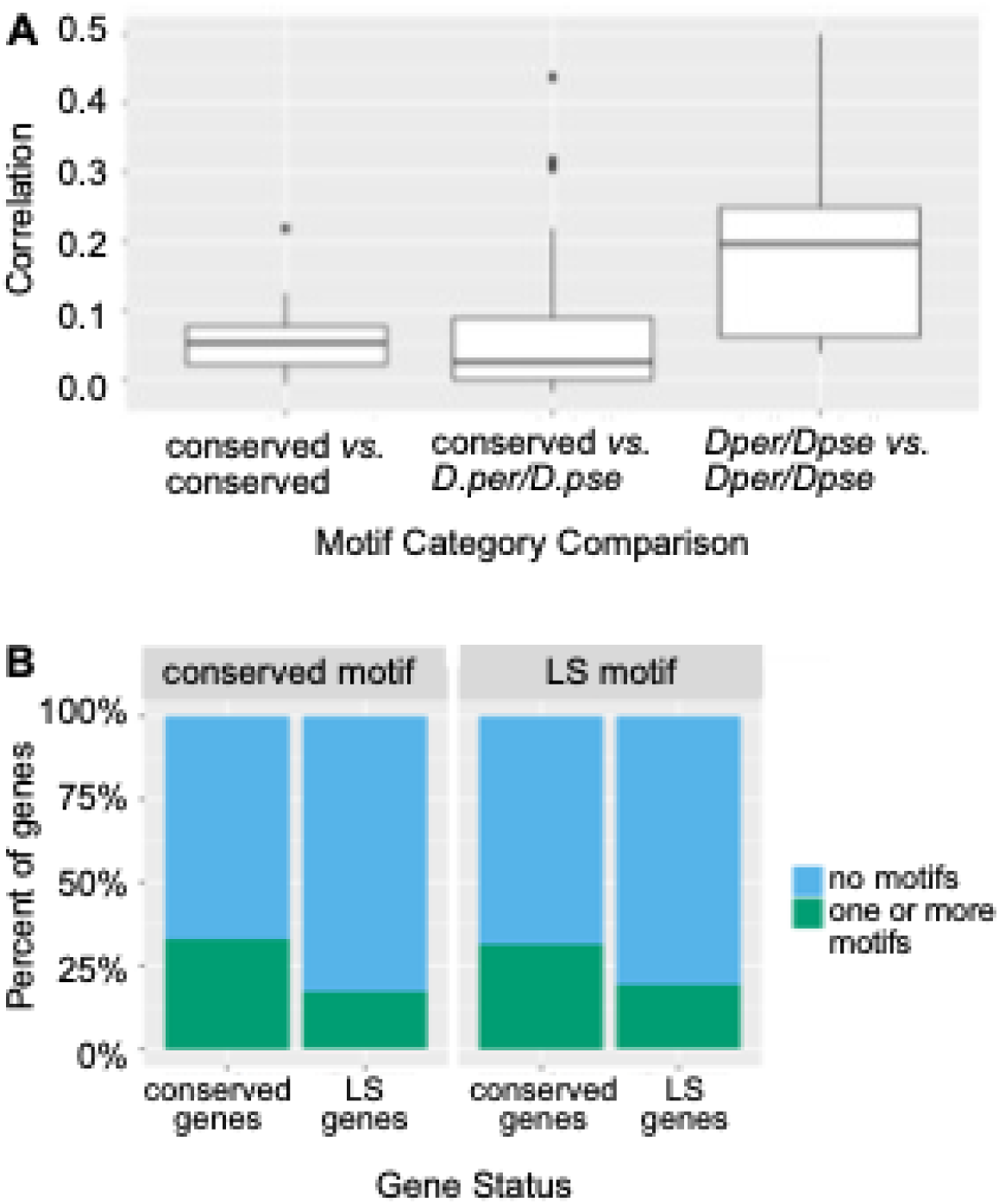
Novel maternal genes are not associated with novel maternal motifs. (A) Correlation coefficient distribution between different sets of motifs. Motifs were classified as conserved if they appeared in most or all species, and species-specific if they appeared in only a single species or lineage (here focused on *D. pseudoobscura* and *D. persimilis* as this is the lineage with a large number of novel motifs associated with maternal transcript expression). Each motif was then compared to each other motif, and correlation coefficients were calculated based on the number of each motif in each gene regulatory region. Conserved motifs have a slight positive correlation, indicating that they appear in regulatory regions of similar sets of genes. Conserved versus non-conserved motifs have a similar distribution of correlation coefficients, indicating that they are intermixed at approximately the same rate. Lineage-specific motifs, meaning those specific to *D. pseudoobscura* (*D. pse*) and *D. persimilis* (*D. per)* have a high correlation with each other, indicating that they appear in a similar set of genes. (B) Conserved genes have more conserved motifs than the *D.per/D.pse* lineage-specific (LS) motifs. Genes that are not conserved across species (LS genes) also show the same pattern of containing more conserved motifs than non-conserved motifs.

Given that there are a number of transcripts that are only maternally deposited in *D. persimilis* and *D. pseudoobscura,* and that we observe several maternal motifs that are specific to these species, we wanted to explore the relationship between these two phenomena. Specifically, we hypothesized the regulatory changes reflected in the species-specific motifs might be responsible for the new maternally deposited transcripts present in these species, and thus species-specific maternally deposited genes might be enriched for species-specific motifs. By examining the motif content of regulatory regions of genes based on the specificity of both genes and motifs, we found this not to be the case (Figure 3B). Instead, the motifs that are common among all *Drosophila* are more common in the *D. persimilis/D. pseudoobscura-*specific genes. Therefore, while there is lineage-specific maternal regulation and lineage-specific maternal deposition in these species, the former is not primarily responsible for the latter.

### Lineage-specific transposable elements are enriched in *D. persimilis/D. pseudoobscura* novel motifs

The abundance of species-specific motifs in the *D. persimilis* and *D. pseudoobscura* genomes is striking because there are considerably fewer differences in motifs between the species sampled with the largest divergence times. Given the rapid proliferation of these motifs, we hypothesized that transposable elements (TEs) could be a vehicle for motif proliferation.

To evaluate whether TEs could be responsible for the propagation of *D. persimilis/D. pseudoobscura*-specific motifs in these species, we tested whether the motifs were enriched in TEs unique to these species. We used the TEs described in a previous study (Hill and Betancourt 2018) which were identified as being specific to the *Drosophila pseudoobscura* group, an observation that we were able to independently verify (Figure S1). We used BLAST to determine the TE locations in all of our reference genomes (see Methods). We calculated enrichment of motifs within these TEs by comparing the number of motifs that occurred within the TE bounds given by BLAST coordinates to the expected number based on the overall length of the TE. Approximately 10.2% of *D. persimilis* and *D. pseudoobscura-specific* TE/motif combinations showed significant enrichment after Bonferroni correction (Figure 4B). In contrast, none of the conserved motifs were enriched in the *D. persimilis/D. pseudoobscura* TEs after Bonferroni correction. We observed a similar enrichment of *D. melanogaster* motifs in *D. melanogaster* TEs, although to much lesser extent (Figure 4A). TEs could therefore be an explanatory mechanism by which motifs regulating maternal transcription could propagate throughout these genomes. This is consistent with other cases, such as with a species-specific TE propagating binding sites for the MSL-mediated dosage compensation complex on the neo-X chromosome of *D. miranda* (Ellison and Bachtrog 2013), thus facilitating the rapid gain of dosage compensation on a new X chromosome in this species.

**Fig 4:**
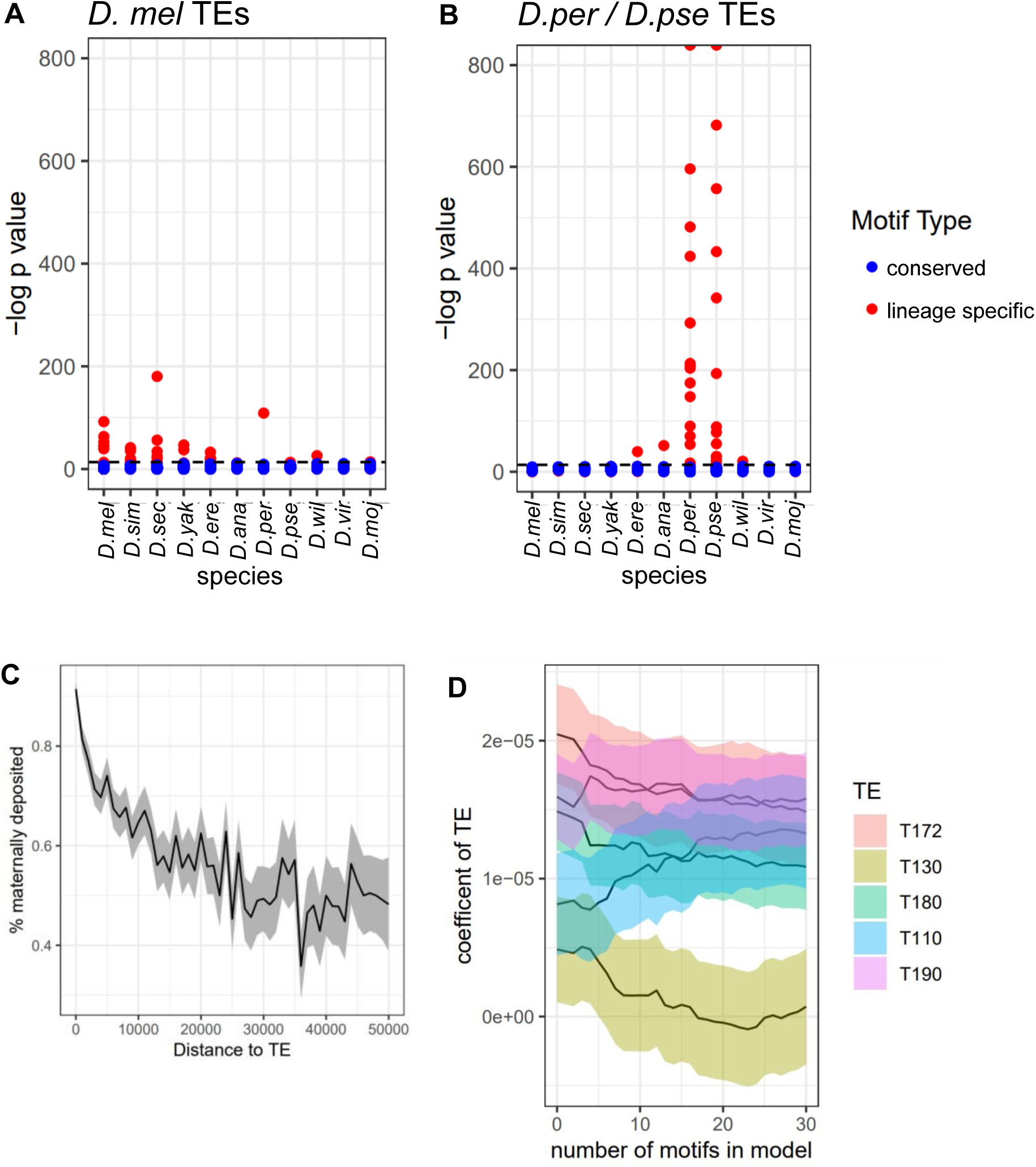
Species-specific motifs are associated with TEs. (A, B) Enrichment of motifs within TEs. Each point represents a single TE-motif pair. Motifs are colored blue to indicate that they are enriched across species, or red to indicate enrichment in a single species or group. (A) Enrichment of motifs within *D. melanogaster* TEs. Due to differences in data sources, the TEs labeled as *D. melanogaster* were discovered in, but are not necessarily exclusive to *D. melanogaster,* and may occur in other species. The conserved motifs are generally not enriched in these TEs, but some species-specific motifs are enriched. (B) Enrichment of motifs within TEs that are known to be *D. persimilis/D. pseudoobscura*-specific. Conserved motifs are generally not enriched within these TEs, but the *D. persimilis/D. pseudoobscura* specific motifs are highly enriched in *D. persimilis and D. pseudoobscura*. *D. persimilis/D. pseudoobscura* TEs were filtered to exclude TEs that appear frequently in other species. (C) Odds of maternal deposition versus proximity to representative TE T130_X. Proximity to this TE is associated with higher rates of maternal deposition. (D) Relationship of *D. persimilis/D. pseudoobscura* TE proximity to maternal deposition rates with respect to the number of motifs. TEs contain motifs that we observe to increase the odds of maternal deposition, but the increase in maternal deposition odds that we observe exceeds our expectations based on those motifs alone. For all motifs pictured here (see legend) except T130, adding additional motifs to the model does not completely explain the correlation with increased transcript abundance with the TE.

Next, we asked whether there was a direct relationship between these *D. persimilis/D. pseudoobscura* TEs and maternal gene expression. When examining transcript abundance measurements of genes that are physically close to these TEs, we find that proximity to some of the *D. persimilis/D. pseudoobscura*-specific TEs is correlated to maternal transcript levels (Figure 4C). It is possible that this is due to the motifs themselves or a consequence of the bias for TEs to transpose into transcriptionally active regions. To untangle some of these potential effects, we employed linear modeling to gauge the effect of both motif content and TE proximity on gene expression. We observe that many TEs are positively associated with maternal transcript levels even when accounting for TE content (Figure 4D), implying that factors within some TEs are contributing to maternal transcript abundance or that these TEs are simply propagated into already active regions at a higher rate.

### Conserved motifs interact with additional factors in a species-specific manner

In addition to motif content, we examined several other factors that contribute to gene expression, such as gene length, orientation, and expression levels of adjacent genes. Some of these factors have species-specific interactions with motif content that could help explain how regulation changes across species in early development. We used generalized linear models to identify the most important factors in terms of species-specificity. The most significant effects for maternal genes are motif interactions with the distance to the nearest upstream gene at stage 2 (Figure S2A), with an example of a motif displaying this interaction in Figure 5A. At this stage, nearby genes tend to be co-expressed (Omura and Lott 2020) so it is unsurprising that gene distance might be important, but we do not yet understand why these motifs would behave differently relative to gene distance in different species (when species-specific gene spacing is accounted for). For zygotic genes (stage 5), the most important species-specific factor is the interactions between motifs and maternal (stage 2) expression level (Figure S2B). This interaction between maternal (stage 2) transcript level and motif content at stage 5 is particularly striking (Figure 5B,C), as one of the motifs that display this behavior is the motif bound by GAGA factor, a well-studied regulator in early development (Adkins, Hagerman, and Georgel 2006; Li and Gilmour 2013; Tsai et al. 2016). While GAGA factor is generally understood to increase expression (Adkins, Hagerman, and Georgel 2006), we find that this effect is not consistent throughout *Drosophila*. The GAGA motif is associated with the activation of genes that are not maternally deposited, especially within the melanogaster subgroup (Figure 5B). However, outside of *D. melanogaster*, *D. simulans*, and *D. sechellia*, we observe that this motif is associated with a decrease in expression in genes that were maternally deposited. It is possible that this repressive action is due to an interaction with *lolal.* GAGA is known to recruit lolal and is thought to mediate polychrome group complex formation (Mishra et al. 2003). *lolal*, in turn, itself has species-specific transcript levels that are correlated (regression p<0.05, see Methods, Figure 5D) with the apparent repressive action of GAGA.

**Fig 5:**
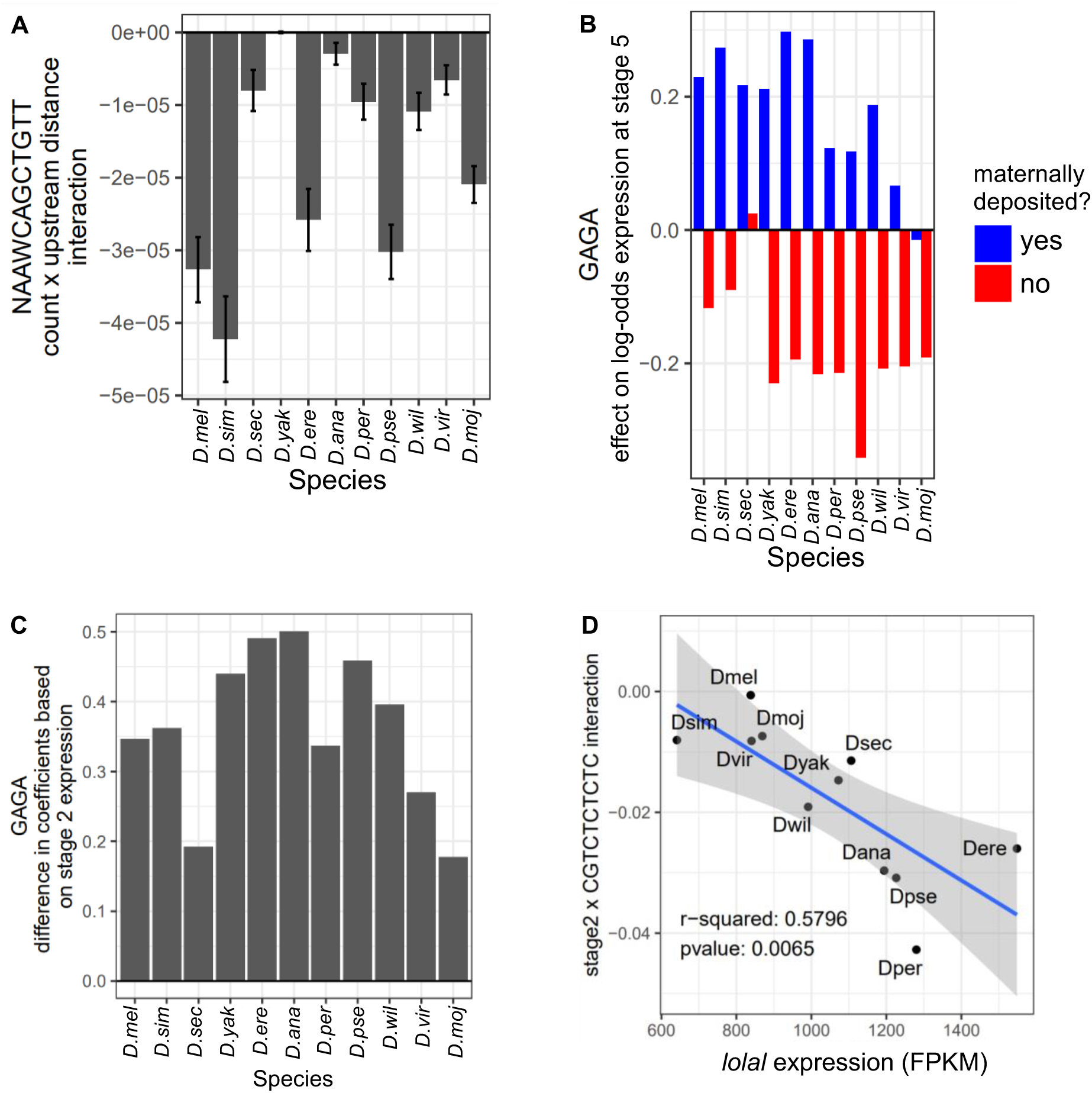
Motifs have species-specific interactions with other factors. (A) An example of a motif displaying species-specific interactions between motif count and upstream gene further away, taking into account the species-specific gene spacing. (B) Effects of GAGA motif on stage 5 expression depend both on species and whether transcripts are maternally deposited. (C) The difference in the effect of these two conditions is also species-specific, with smaller differences observed in *D. melanogaster, D. simulans, D. sechellia, D. virilis*, and *D. mojavensis.* (D) These differences in effect on log-odds expression are correlated with *lolal* transcript abundance. As lolal is associated with the repressive action of GAGA, it is possible this represents a *trans* factor conferring species-specific effects to this motif.

### GO terms associated with the same maternal motifs differ in different species

Given we found conserved motifs to have slightly different behavior in different species, we asked whether conserved maternal motifs (Omura and Lott 2020) could be regulating genes with different functions in different species. To address this, we performed GO analysis on the set of genes associated with each motif in each species. To measure similarity, we then used Pearson correlation on the log p-values (see Methods). This approach is somewhat limited by the *D. melanogaster*-based GO annotations, as genes are assigned GO categories based on their *D. melanogaster* orthologs, potentially leading to bias. We found that different motifs had different levels of similarity in GO category association between species. Overall, gene annotation similarity was highest among members of the *D. melanogaster* subgroup (Figure 6), and lower outside of this lineage. This suggests that even though motifs are conserved they could be regulating different sets of genes in the most diverged species. Interestingly, *D. sechellia* displayed a lower-than-expected similarity to its closest relatives, a finding that requires further exploration beyond the scope of this study.

**Fig 6:**
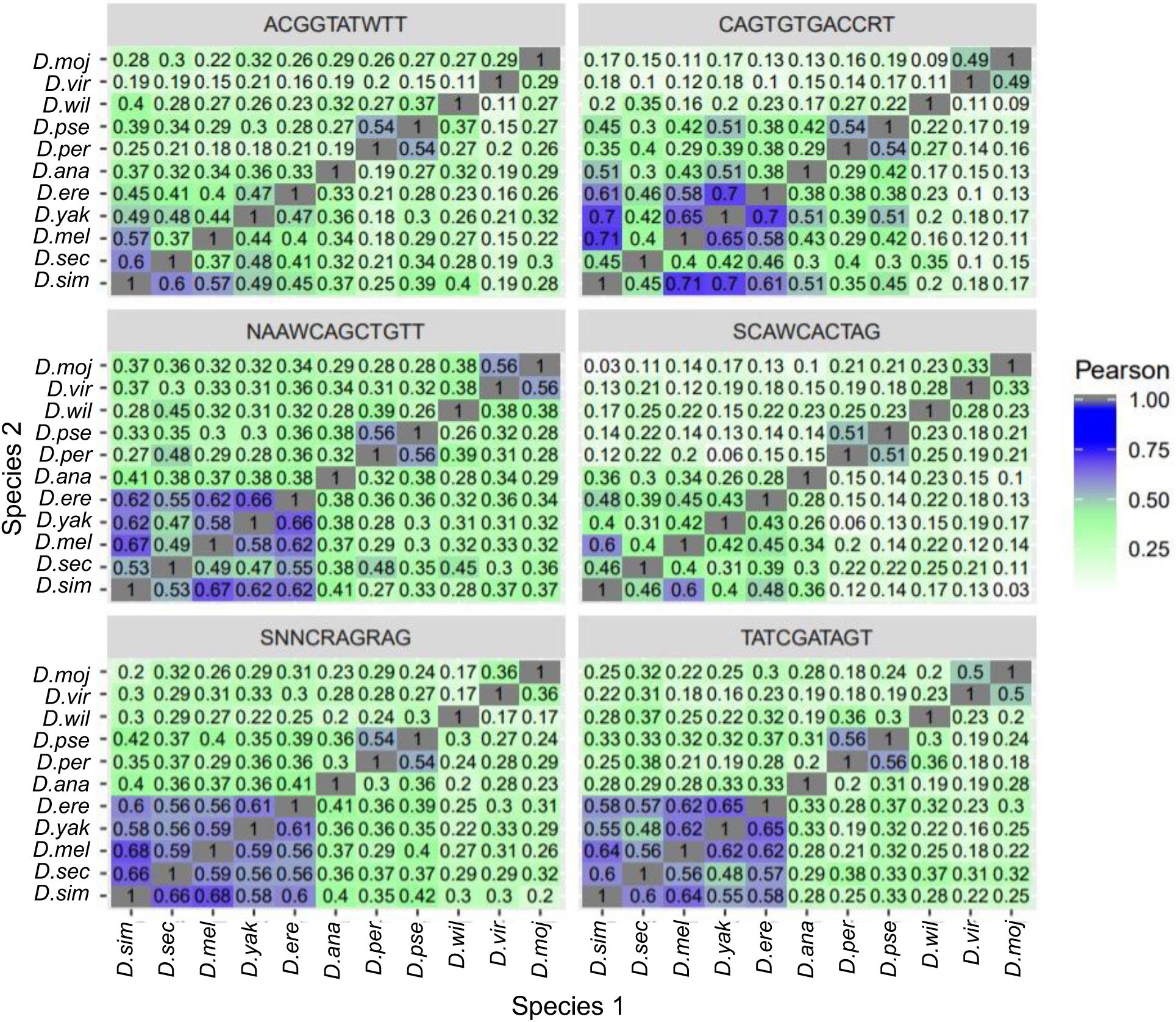
GO terms associated with different motifs have different levels of conservation. In order to determine the functional similarity between genes associated with specific motifs across species, eight common maternal stage motifs were selected for GO analysis. Functional GO term enrichment was calculated for the targets of each motif, and Pearson correlations between GO categories were determined for between each pair of species. Despite GO annotations being based off of *D. melanogaster*, we observe cases where distantly related pairs of species, (for example. *D.mojavensis* and *D.virilis* for the motif TATCGATAGT) display higher correlation than other similar comparisons.

### Given the motif content predicted gene expression differs between maternal and zygotic genes

Given our knowledge of how various factors affect maternal or zygotic transcript levels, we next asked whether this information could be used to explain differences in transcript abundance across species. Thus we compared pairs of orthologues from each species in terms of motif content and expression difference (Figure 7) at each stage. Given distinct transcript abundance values between genes that are and are not maternally deposited (Omura and Lott 2020), and the abundant shared motifs at the maternal stage, we expected to find similar motif content in similarly expressed genes. At both stage 2 and stage 5, we found that if a pair of orthologs were both lowly expressed, they tended to have similar motif contents, while pairs of orthologs that were both highly expressed tended to have quite different motif contents (Figure 7). In the cases where a transcript was present in one species but its ortholog was not present in another, their motif contents interestingly tended to be slightly more similar than pairs that were both present. This indicates that motifs play a lesser role in determining whether a gene is off or on than perhaps expected. The strikingly different motif content in pairs of expressed orthologs likely speaks to the selection pressures on transcript levels: as long as transcript levels are maintained, the combination of motifs that facilitate this level of expression can evolve. At stage 2, within pairs of genes that are both expressed, there is a visible trend for pairs with similar motif contents to have more similar expression levels. This suggests that changes in motif content could result in differences of expression level, but not necessarily to the point of turning genes off or on. In contrast, at the zygotic stage, we see no such trend and the motif content distances between expressed genes appears more random. This suggests that at the maternal stage, motif content may serve a more quantitative role in determining transcript abundance, while at the zygotic stage motif content may play a more qualitative role.

**Fig 7:**
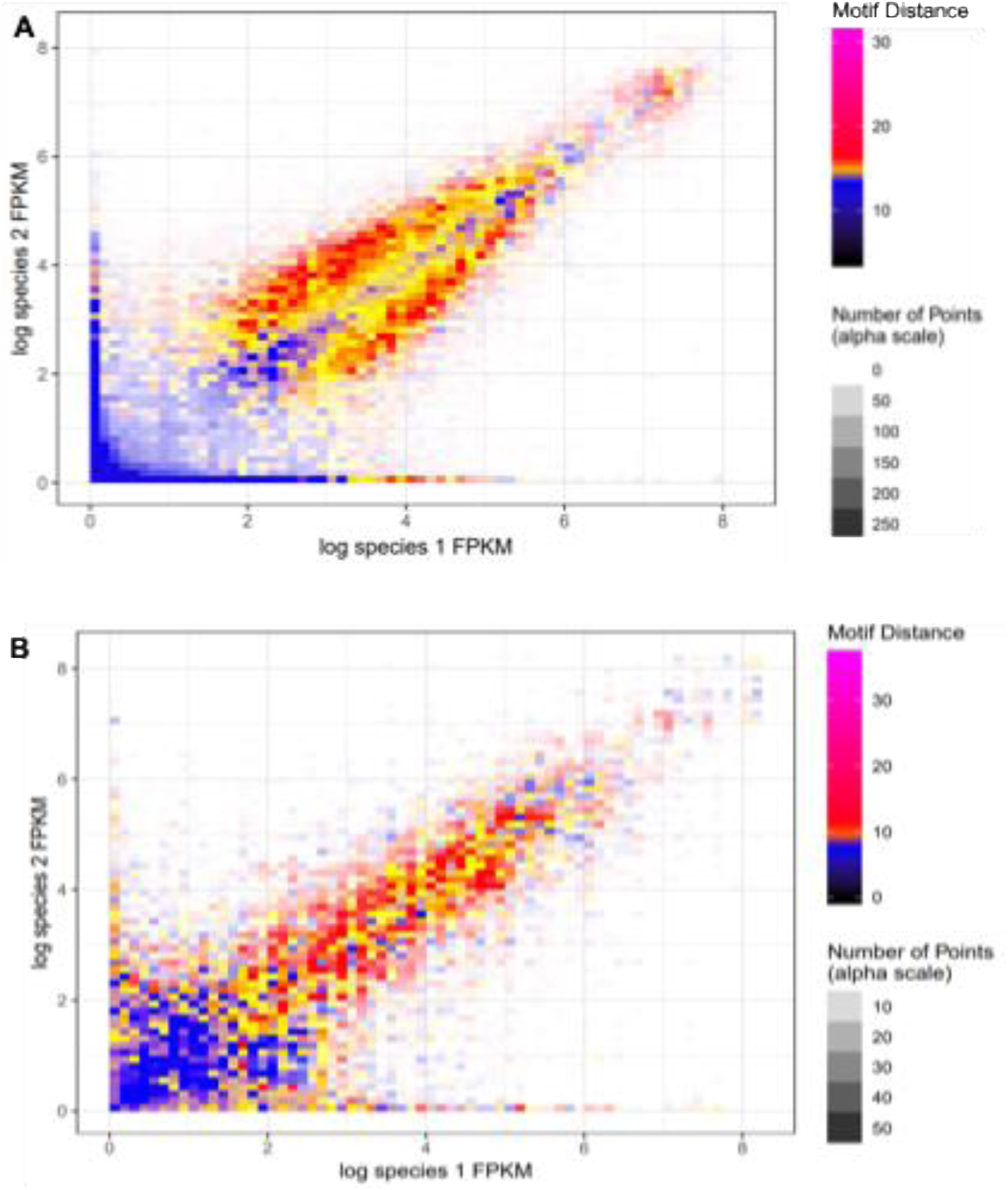
Motif analysis is insufficient to explain transcript abundance. To explore the relationship between transcript level changes and motif differences, each pairwise combination of orthologs for each gene is analyzed. The x-and y-coordinate correspond to the transcript levels of each comparison. Comparisons are binned, and each bin is colored with hue corresponding to the difference in motif content, here described as “motif distance” (see Methods). The color yellow is the median change in motif content and alpha corresponds to the number of representative comparisons. A) In Stage 2, many pairs of orthologues where at least one species has low expression have similar motif contents (blue cloud in the lower left corner). Many pairs of orthologs where both species have higher expression levels tend to have more different motif contents (red and yellow diagonal), though pairs with very similar expression levels between species can have similar motif contents (blue diagonal). B) In stage 5, ortholog pairs where one gene is expressed while the other is not are more likely to have different motif contents (red and yellow along the x and y axis), but again many pairs of orthologues where both species have higher expression levels still often have different motif contents (red and yellow diagonal).

Rather than correlating transcript level differences with motif changes, we can also try to predict transcript levels based on motif content and other factors. To achieve this, we used logistic regression implemented in R (R Core Team 2014) to evaluate the probability of a gene being expressed in either stage (see Methods). Utilizing the motif counts, species-specific motif interactions, and other parameters discussed previously in Figure S2, we were able to classify genes as either expressed or not expressed with an AUC = 0.8733 and 0.7792 at stage 2 and stage 5 respectively. This model is likely overfitted due to the large number of possibly non-functional motifs, yet still produces a suboptimal result. This indicates that there are additional factors that play an important role in gene expression we are not accounting for, or that the factors we are measuring do not behave in a way that is suitable for simple logistic regression.

Because logistic regression may fail to capture the biological complexity of our motifs, we implemented random forests to evaluate the relative importance of different contributing factors to early gene expression. At the maternal stage, we find that the most important factors are the proximity and expression of adjacent genes (Figure 8A). We also find that the most prominent motifs, such as DREF, have the most importance. This is consistent without our observation that chromatin architecture is likely a key factor in transcriptional regulation at this stage (Omura and Lott 2020). At the zygotic stage, while we again find a strong emphasis on some chromatin related factors (Figure 8B), we also see a difference in motif importance. For zygotic genes, we find a less centralized concentration of motif importance, with more, weaker motifs displaying a more uniform importance (Figure 8C). This is consistent with our understanding that spatiotemporally specific motifs are acting independently on smaller subsets of genes at this stage. Interestingly, multiple factors that one would expect to be most closely related to stage 2 are rated highly at stage 5, including residual (<1 FPKM) expression at stage 2. This is surprising, considering the transcripts at the two stages come from distinct genomes from different life stages (that of the mother and the zygote). It is feasible that gene accessibility may be correlated between the maternal and zygotic genomes, and therefore even very low expression in one could be indicative of the potential for higher expression in the other.

**Fig 8:**
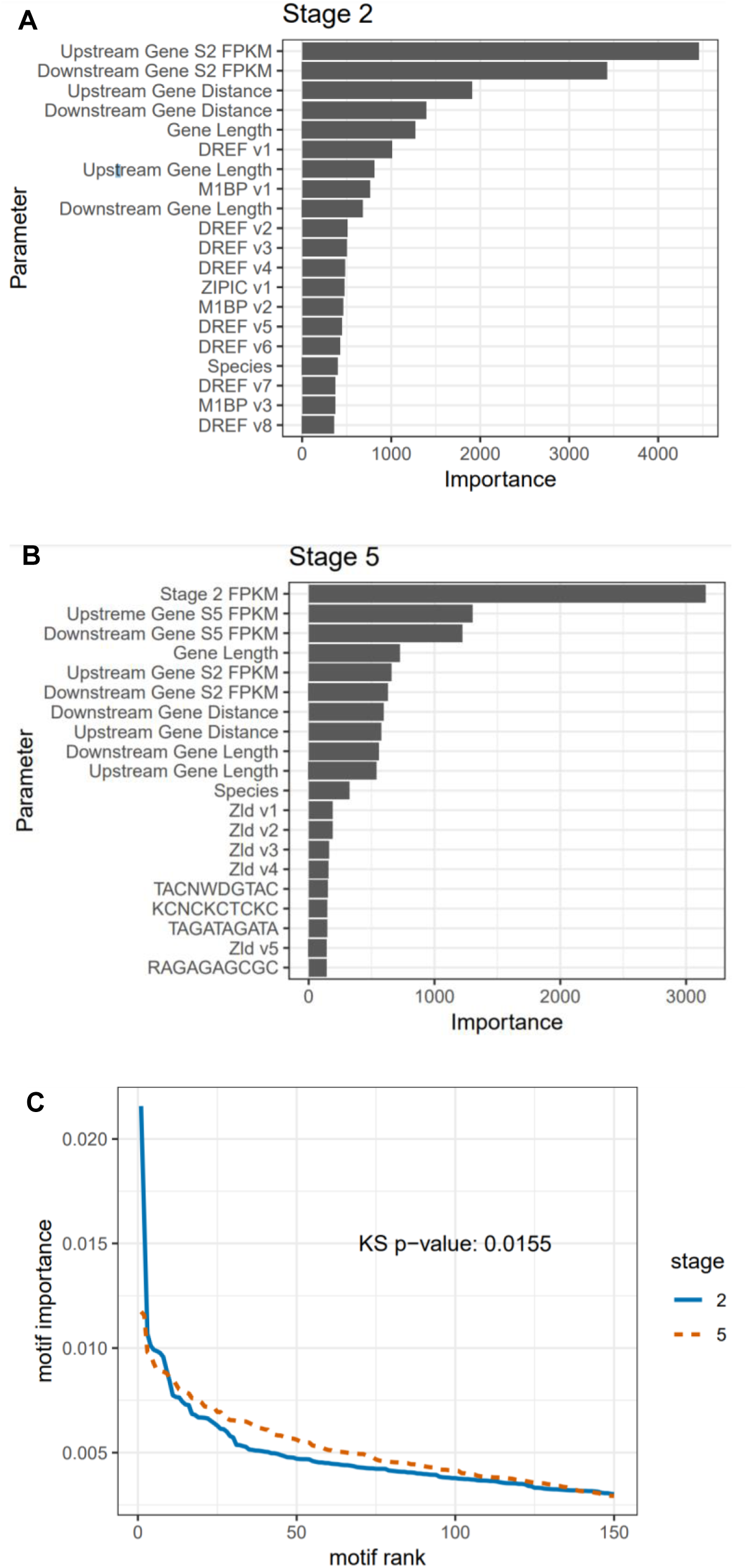
Specific motifs are less predictive of transcript abundance than chromatin. (A) Random forest-derived importance scores for each factor in maternal deposition. (B) Random forest-derived importance scores for each factor in expression at stage 5. (C) Importance of the top 150 motifs at each stage. The y-axis represents the importance scaled to the model total.

### Maternal genes that change in expression between species are more likely outside the co-expressed clusters

We examined maternal transcripts whose abundance changed between species to gain insight into how these changes may evolve. We did not measure this phenomenon for zygotic genes due to complications involving the definition of consecutively expressed genes at this stage when any transcripts measured could be either maternal or zygotic in origin. For maternal genes, we found that genes with different expression patterns across species tend to be more physically isolated in the genome in terms of consecutive co-expressed genes (Figure 9), with the most likely genes to change expression being flanked on both sides by non-expressed genes. This suggests that the mechanisms that are responsible for maintaining expression for multiple consecutive genes are more robust than those controlling single genes. This is consistent with control of maternal gene expression at the level of chromatin state, as it may be difficult to activate transcription of a single gene in a region of closed chromatin or repress transcription of a single gene in an open and heavily transcribed region.

**Fig 9:**
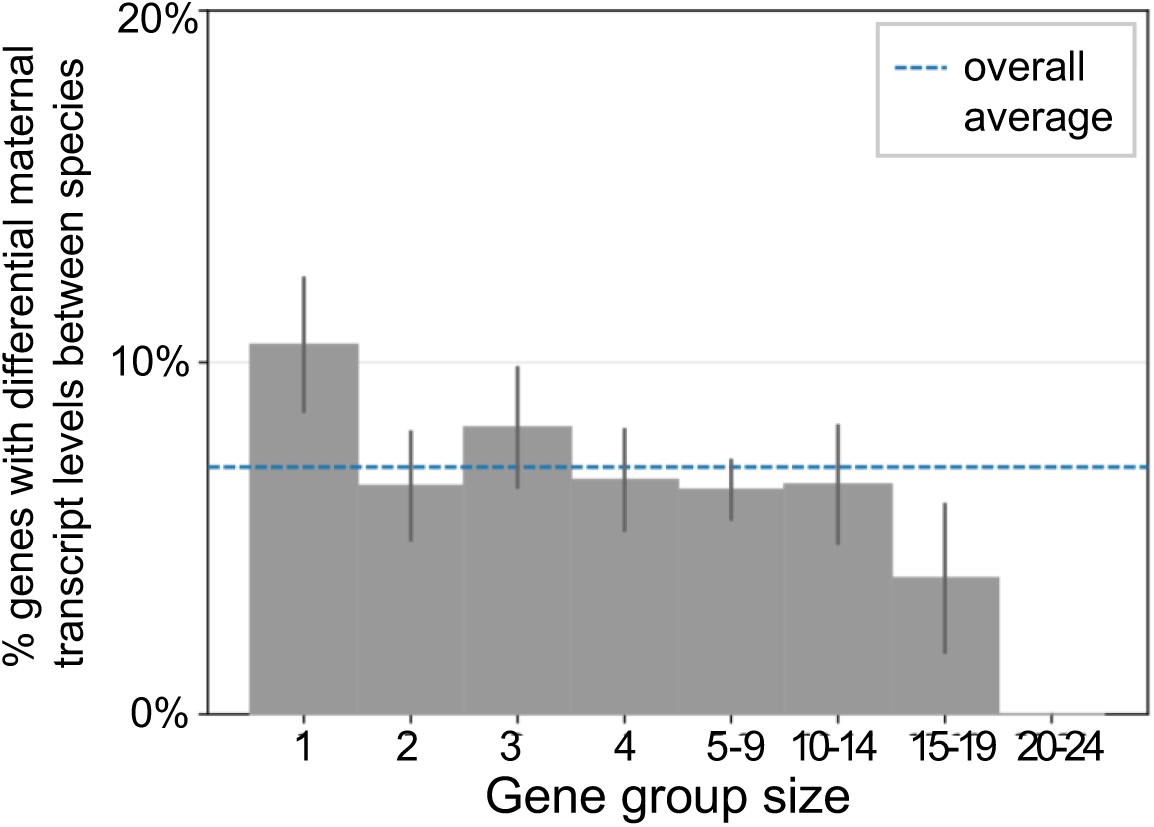
Physically isolated genes are more likely to have differences in transcript level between species. Given the possibility that maternal gene regulation appears to occur on the level of chromatin topography, genes that are physically isolated along the chromosome are a good candidate for observing regulation across species. Representative species *D. simulans* was chosen for its average behavior. Gene clusters are here defined as genes that are expressed and adjacent on the chromosome. Group size is the number of genes within the cluster. Error bars represent 95% confidence intervals. The dotted blue line represents the average changing gene rate disregarding gene groupings. Genes with group size 1, i.e. flanked by genes that are not deposited, are more likely to change their expression between species.

## Discussion

The early stages of development are well conserved across *Drosophila*, even at the level of transcript abundance of individual genes (Atallah and Lott 2018). This would suggest high levels of developmental or evolutionary constraint to keep transcript levels in a similar range. Yet, evolution of maternal and zygotic transcripts does occur, as there are some observable differences in transcript presence and levels across species (Atallah and Lott 2018; Heyn et al. 2014). Here, we wanted to ask how evolution of transcript level regulation occurs for maternal and zygotic transcripts in the early *Drosophila* embryo.

### Evolution of transcript levels through lineage-specific regulators

One possible explanation for how new transcript representation or transcript levels can evolve is through the evolution of regulatory proteins. These regulators can be produced from genes which were not transcribed at a particular stage, or if present, did not previously have a role at that stage. Alternatively, new regulators could be products of novel genes, or could represent significant functional changes in existing proteins. In our study, all of these potential types of changes in regulatory proteins would appear as lineage-specific regulators.

At the maternal stage, the evolution of lineage-specific regulation appears to be a rare event, with the most prominent examples of this coming from *D. persimilis* and *D. pseudoobscura.* In these species, propagation of TEs may have facilitated the emergence of motifs for lineage-specific regulators, in a similar manner to the propagation of binding sites for the X chromosomal dosage compensation complex on the neo-X chromosome of *D. miranda* (Ellison and Bachtrog 2013). While these species do have a greater number of evolved differences in gene expression than expected given their position on the phylogeny (Atallah and Lott 2018), this is not due only to new regulators, as lineage-specific genes (those identified as present or absent only in this lineage) are not enriched in lineage-specific motifs. At the maternal stage, where we believe regulation of transcription occurs on the broader level of chromatin state and topologically associated domains (TADs, regions of highly interacting DNA that are separate from other such regions) rather than individual genes (Omura and Lott 2020), it may be more difficult for gene expression to be modified through individual binding sites, although this is still possible. It may also simply be more difficult at this stage to activate transcription of a gene in a region of closed chromatin or repress transcription in a highly transcribed region. Given the importance of factors such as gene distance, length, and strand, it is possible that individual changes in gene expression are brought about by changes in these factors. However, it seems most likely that evolved changes in transcript abundance at the maternal stage may be due to changes in abundance of the chromatin-modifying regulatory proteins. Previous work indicates that many changes in maternal gene expression between closely related species are due to trans-regulatory changes (such as those in regulatory proteins), and that closely related species also have differences in the abundance of transcripts for the important chromatin regulators of maternal transcription (Cartwright and Lott 2019).

In contrast, evolution of lineage specific motifs occurs more frequently during the zygotic stage, where we also observe more differences in gene expression. Whether these lineage-specific regulators are responsible for lineage-specific expression is unclear. But it is more plausible for zygotic genes than with maternal transcript regulation given the much larger number of regulators at this stage, presumably regulating smaller numbers of genes in more spatiotemporally precise ways. One approach to address what drives changes in transcript representation or transcript abundance for specific genes is by directly analyzing regulation of this subset of genes that change, but we lacked the statistical power to do so in this study.

### Evolution of transcript levels through changes in existing regulators

Another possible mechanism for how transcription can evolve at the maternal or zygotic stages is through evolving changes in the manner in which conserved regulators act. One example of this we observe is that at both stages, we identify cases where the same motif appears to have differing effects on the gene it is likely regulating depending on the species. Although we don’t yet know the mechanistic underpinnings of this phenomenon, our ability to measure this effect is an indication that it occurs with some frequency over evolutionary time. GAGA factor, a regulator well-studied for its effects on zygotic transcription (Adkins, Hagerman, and Georgel 2006; Li and Gilmour 2013; Tsai et al. 2016), is an example of a regulator that has differing effects in different species. Because this was discovered using methodology that examines genome-wide trends rather than specific binding sites, we conclude that this effect is likely to affect a large number, perhaps all, of genes targeted by GAGA. Like with lineage-specific regulators, changes in conserved regulators that affect expression of a large number of genes seem like they would be selected against, as it would likely be detrimental. However, as with our examples of gains of lineage-specific regulators, we observe this occurring with enough frequency to be detected by our methods. Indeed, the changes in the signals of how conserved regulators act across species suggests that even conserved motifs could gradually decrease their effect and potentially be replaced in distantly related species, suggesting a mechanism for turnover of conserved regulators over longer periods of evolutionary time. Additionally, we found a number of conserved motifs that show species-specific interactions with gene spacing at the maternal stage, and with maternal transcript level at the stage after zygotic genome activation. While these also represent changes that impact a large number of genes, the fact that they rely on an additional element means that this number is a subset of genes controlled by a particular regulator, and therefore may evolve changes without affecting all targets of this regulator. Thus this additional specificity might mean these species-specific changes are less deleterious overall.

### Evolution of transcript levels through changes in motif binding sites

Transcription can also evolve through changes in binding sites or larger scale reordering of regulatory sequence at individual genes. Given that maternal regulation is likely occurring at the level of chromatin state, this seems less likely to globally affect transcription at the maternal stage. As we had previously found that individual maternal genes did not show conservation of the same set of binding sites for chromatin modifiers (Omura and Lott 2020), it seems plausible that some of these regulators may be redundant, allowing for compensatory evolution of binding sites without the loss of expression. On the other hand, changes in regulatory sequence and the associated motifs seem more likely to be a mechanism through which zygotic transcript levels can evolve. This mechanism would be consistent with what is known about how regulation can occur at this stage, through precise regulation at individual genes in time and space (Jaeger, Manu, and Reinitz 2012; Briscoe and Small 2015). The regulatory regions that have been well-studied for particular zygotic genes show that expression depends to some degree on combinations of particular transcription factor binding sites in certain numbers and arrangements, in individual regulatory regions (Arnosti et al. 1996). However, previous research also shows that there can be considerable rearrangement of regulatory DNA over evolutionary time without changes in expression of these precisely regulated genes (Michael Z. Ludwig et al. 2005; M. Z. Ludwig, Patel, and Kreitman 1998), despite the observation that changes in regulatory DNA can be responsible for changes in transcript level. Additionally, in a previous study in closely related species, changes in both cis- and trans-regulatory factors were identified as important to the evolution of differences in transcript level for zygotic genes at the stage also examined here (Cartwright and Lott 2020). While that study was not designed to look at either specific binding sites or changes in specific regulators, it seems likely that changes in both may be important to evolving transcript level differences for zygotic genes.

### Selection and post-transcriptional regulation

While our primary focus has been on transcript abundance, selection ultimately acts on phenotypes, which we expect to be most directly dependent on protein abundance. This seems especially important to consider at the maternal stage, where there is considerable post-transcriptional regulation. Since transcript abundance itself cannot be changed by differential transcription before the zygotic genome is activated, it has been demonstrated that post-transcriptional regulation is especially important for maternal transcripts. Maternal transcripts are regulated at the level of stability (Cui et al. 2013; Eichhorn et al. 2016), are subjected to programmed degradation (Tadros et al. 2007; Bashirullah, Cooperstock, and Lipshitz 2001), and are in some cases specifically localized within the oocyte and later the embryo (Medioni, Mowry, and Besse 2012; Vazquez-Pianzola et al. 2017). These mechanisms of post-transcriptional regulation have been shown to alter the protein levels associated with maternal transcripts (Sallés et al. 1994). However, previous studies indicate an even higher level of conservation of maternal transcript levels overall (Atallah and Lott 2018; Heyn et al. 2014) than zygotic transcript levels. In addition, the fact that maternal transcription involves a large proportion of the maternal genome to be quickly transcribed and that regulation is likely occurring at the level of chromatin would seem to indicate that transcription is controlled by imprecise mechanisms. Perhaps what can reconcile this apparent paradox is if the nature of the imprecise chromatin-level transcriptional control means that selection on protein levels to evolve changes is more effectively accomplished at the post-transcriptional levels of regulation. This would produce the highly conserved pattern of maternal transcripts that we see as a product of a mechanistic constraint, where protein levels could still be fine-tuned by changes in post-transcriptional mechanisms. With fewer post-transcriptional control mechanisms for zygotic transcripts and more opportunities to change expression at the level of transcription, we would expect to see a larger number of changes in transcript levels we see for zygotic genes.

## Conclusions

In this study, we investigated the evolution of gene regulation in the early embryo, and the associated effects on maternal and zygotic transcription. These early developmental stages are critical, and the developmental processes and transcript levels during these stages are highly conserved. Yet some variation in transcript level does occur, so through what mechanism does this variation evolve? We examined regulatory regions of transcribed genes to detect sequence motifs in 11 species of *Drosophila* to explore how changes in motifs relate to changes in gene expression from the maternal and zygotic genomes. We found that while most species use a relatively conserved set of motifs, we also find some examples, particularly in *D. persimilis* and *D. pseudoobscura,* where very different motifs are utilized in addition to the conserved motifs. These new motifs appear to be regulating genes in concert with conserved motifs. We found evidence that transposable elements (TEs) may both have been a vehicle for motif propagation, as well as contribute to expression in their own right. At the zygotic stage, we found that more motifs tended to be species-specific across all the species, without any particular species or lineage showing an overrepresentation of novelty. Additionally, we found that conserved motifs themselves can be used to regulate different sets of genes in different species, at both stages. Furthermore, other species-specific factors can play a role in motif behavior, for example, gene spacing has species-specific interactions with some motifs at the maternal stage. Given the high conservation of transcriptomes at the maternal stage our understanding that chromatin-level regulation with redundant non-specific motifs points to a regulatory environment where evolution occurs via the gradual change of regulator behavior, with some exceptions. In contrast, at the zygotic stage, evolution may be mediated by the relatively rapid turnover of motif binding sites and novel regulators, which is likely responsible for the greater variation in transcript level that we observe at this stage.

## Methods

### Data Acquisition

RNA-seq data utilized for this experiment was generated previously (Atallah and Lott 2018), and is available at NCBI/GEO under accession number GSE112858. GTF files and reference genomes were downloaded from Flybase (Gramates et al. 2017). To determine whether a gene would be labeled as ‘off’ or as ‘on, the overall distribution of FPKMs was analyzed. For all species, for both stage 2 and stage 5, a bimodal distribution appeared, with one peak at 0 and another at ∼e∧3.5 (Figure S3). The commonly used cutoff of FPKM=1 was chosen as it falls between these two distributions.

### Sequence Selection

Preliminary tests were performed to determine which regions were most likely to have regulatory elements. For each gene, several regions were extracted: 10kb upstream, 2kb upstream, 5’ UTR, total introns, total exons,and 3’ UTR. For each region, boundaries were obtained from the GTF and sequences were extracted using bedtools. The 2kb upstream region showed the most distinction between on and off. For the main analysis, UTRs were ignored as not every species had annotated UTRs.

### Motif Discovery

We used Homer (Heinz et al. 2010) to discover motifs in test sets using the background sets as control fastas. Deviations from the default settings include the use of the -fasta flag to specify a custom background file. For stage 2 queries, the test fastas included genes that had a FPKM >= 1 at stage 2 while the control fastas included genes that had an FPKM < 1. For the stage 5 queries, the test fastas contained genes where the stage 5 FPKM >= 1 and the stage 2 FPKM < 1, while the control fastas included genes whose stage 5 FPKM < 1 and stage 2 FPKM < 1. Additionally, we used the -p flag to utilize our computational resources more efficiently. We used -norevopp flag in the case of strand-specific searches and -nlen 10. Motif quality was evaluated based on the Homer-outputted q-values.

### Zygotic Only Motifs

To examine the regulatory elements associated with zygotic genome activation, we defined genes as being zygotic only if they were off at stage 2 and on at stage 5. It is necessary to impose such a restriction as a large (86%) percentage of genes that are maternally deposited are still above the on/off threshold by stage 5, and analysis of their regulatory mechanisms would confound the signal of stage 5 only genes.

### Motif Sharing

To determine whether motifs were shared between species, the Homer-formatted motifs were converted to meme-formatted motifs using chem2meme (Bailey et al. 2009). Tomtom was then used to find matching motifs, using default parameters. For a motif to be considered shared with another species, the Tomtom output threshold of *α* = .05 was used.

### Sequence Similarity

To determine the similarity of genes and proteins, we used Clustal Omega (Chojnacki et al. 2017) multiple sequence alignment.

### Species-specific motif identification

Motifs with different effects in different species were identified using logistic regression. For each motif, we generated two models:

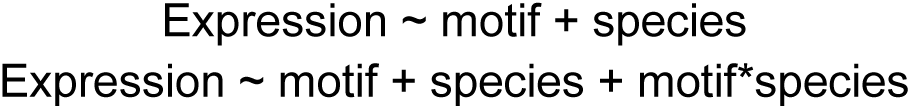

Using the glm function in R using the “stats” package version 3.5.1, using logit as the link function. We used the r logLik function to calculate the log likelihood of each model, and then applied the likelihood ratio test. We used the pchisq function to approximate the statistical significance of adding the interaction terms, and ranked motifs accordingly.

We used a similar technique to find species-specific interaction effects:

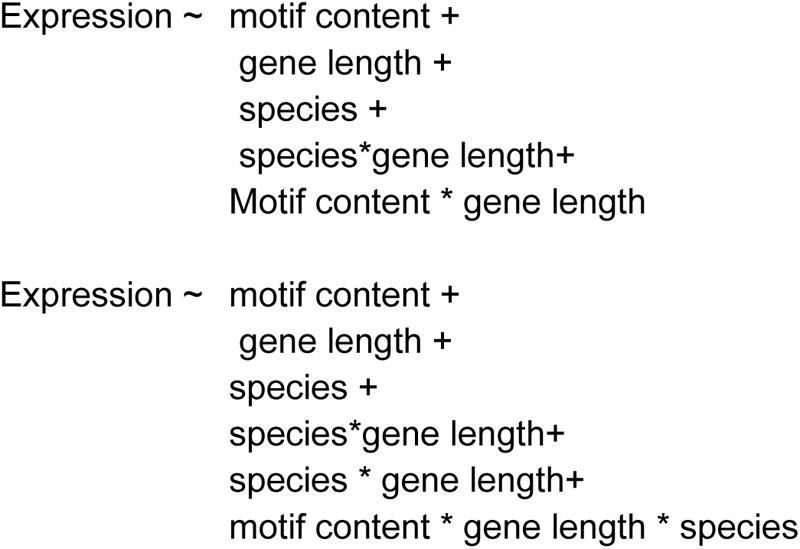

And again comparing the models with the likelihood ratio test. For overall prediction, we calculated AUC using the auc function from the pROC package version 1.14.0.

### Random Forest

ranger(y = as_data[,339], x = as_data[,-339], case.weights=s2_weights, importance = ’impurity’) Random forests were generated using the ranger (version 0.12.1) function in R. Cases were weighted inversely proportionally to the percentage of the expressed genome to account for the differences in expression between stages. Importance was measured using the ‘impurity’ metric.

### Identifying Motifs within TE Boundaries

(Hill and Betancourt 2018) TEs were searched against each reference Genome using Blastn 2.7.1, and the locations were used as TE boundaries. Motifs Locations were identified using the HOMER scanMotifGenomeWide function. Motifs within the TE boundaries were considered to be TE motifs. The total motif lengths were calculated by summing the area within TE boundaries for each TE in each species. Enrichment of motifs within these TEs was calculated by comparing the number of motifs that occured within the TE bounds given by BLAST coordinates to the expected number based on the overall length of the TE. P-values were generated using the poisson distribution.

### GO term similarity

Genes containing conserved motifs were identified in each species, and similarity between sets of genes was determined using TopGO (Alexa and Rahnenfuhrer, n.d.) version 2.34.0. The runTest function was called using the “classic” algorithm and “fisher” statistic for each species. To compare similarities between pairs of species, we used the “cor” function from the “stats” package version 3.5.1.

## Supplemental Figures

**Fig S1:**
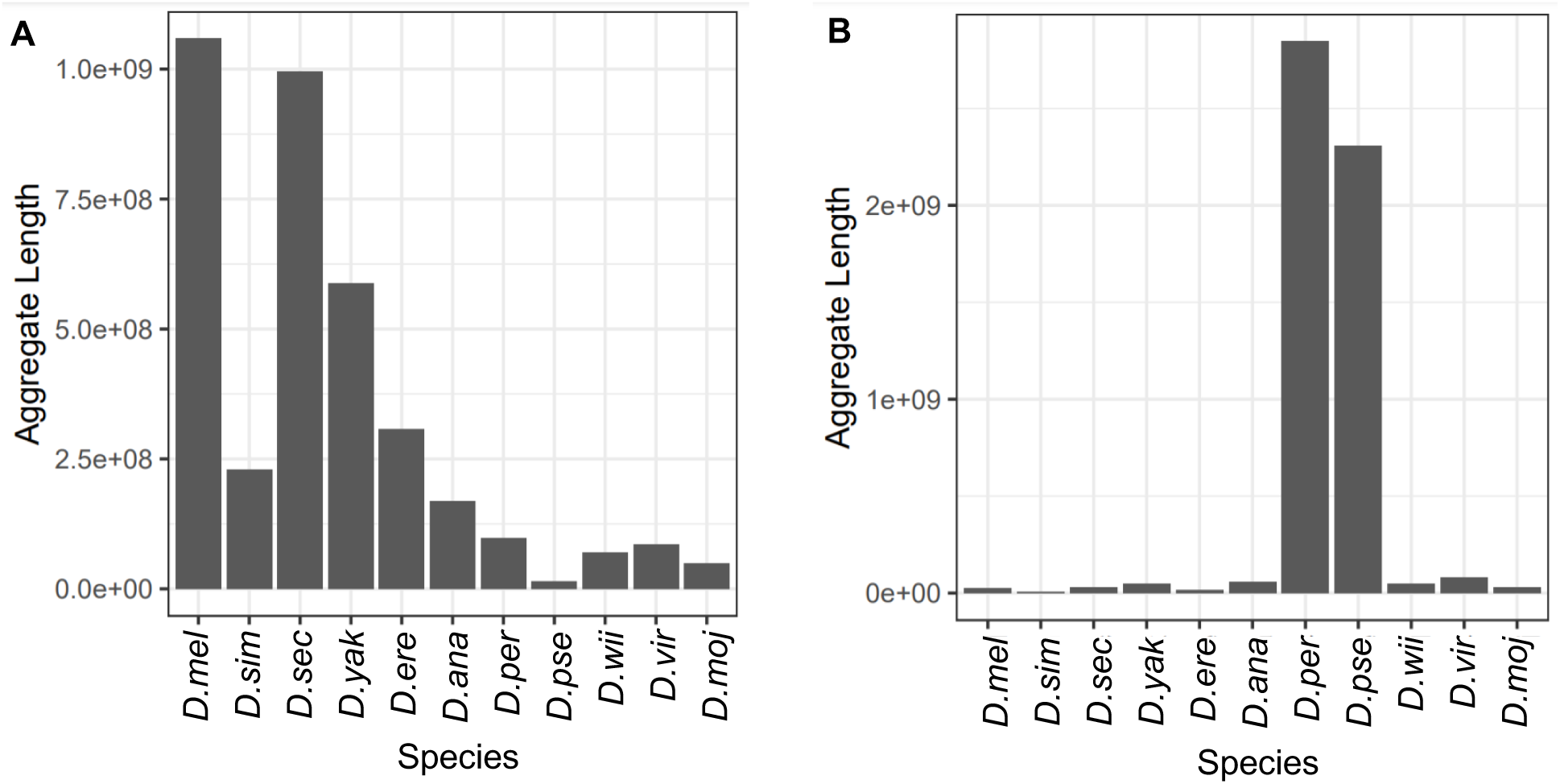
Species-specific TEs are indeed species-specific. In order to determine if our list of D. melanogaster TEs were in fact primarily found in D. melanogaster, we examined the aggregate length of these TEs in each species as a proxy of their abundance. (A) The aggregate length of *D. melanogaster*-specific TEs in different species. The aggregate length is longest in *D.melanogaster,* though other species contain this TE as well. (B) The aggregate length of *D. persimilis/D. pseudoobscura*-specific TEs in each species. The aggregate length is longest in *D. persimilis* and *D. pseudoobscura*.

**Fig S2:**
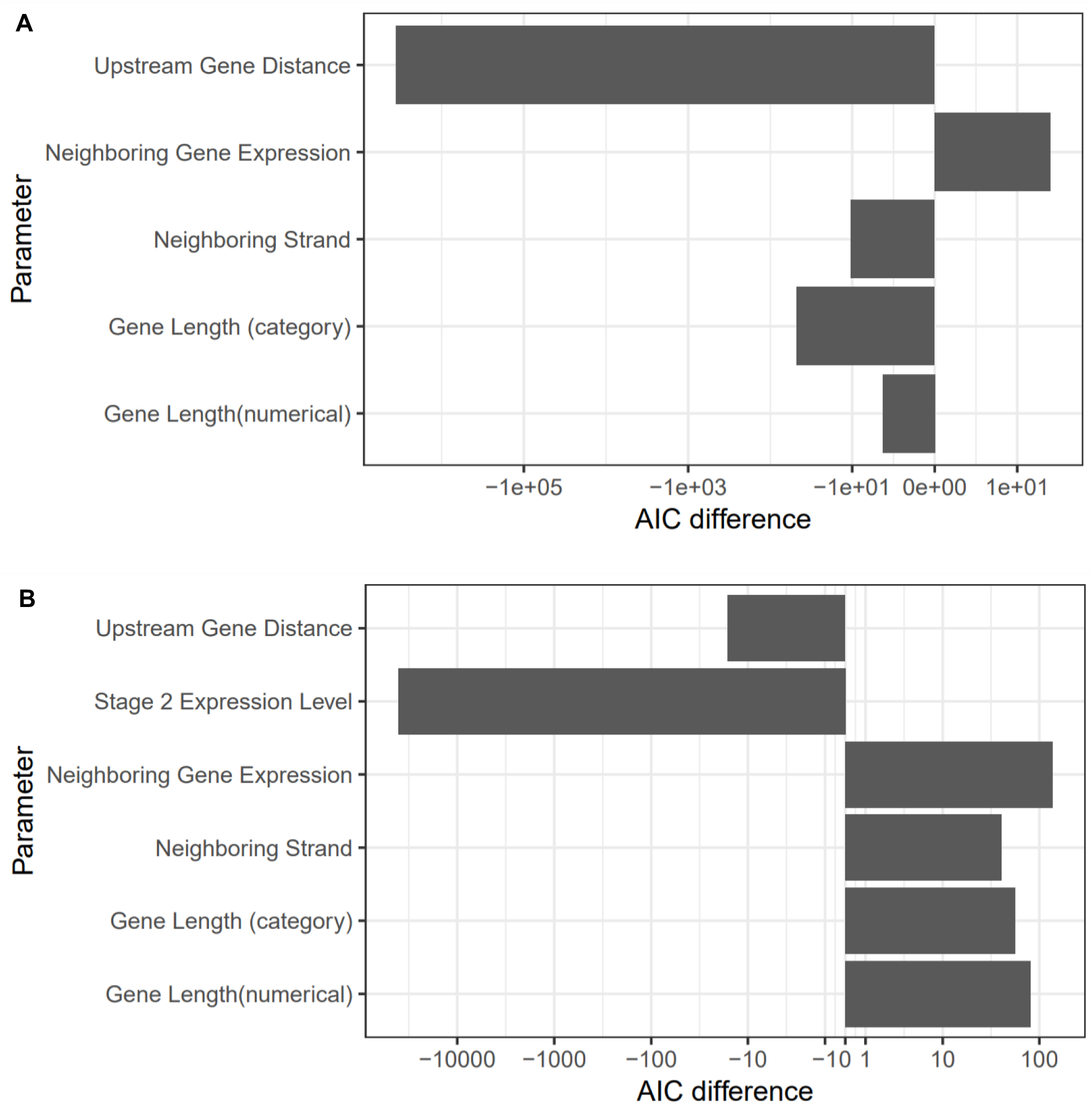
Secondary parameter interaction importance. To search for interactions between motifs content and other factors, we generated generalized linear models with either the interaction term included and excluded, and evaluated them with their AIC. A negative AIC difference indicates that the model with the interaction term performed better, suggesting a possible biological significance to those interactions. Note the pseudo log x-axis. Gene length is measured either as a category (greater or less than 2 kbp) as well as quantitatively. Neighboring gene expression indicates the expression level of the upstream gene, while neighboring strand indicates whether the upstream gene is on the same or opposite strand. (A) At stage 2, upstream gene distance is the only interaction term with a negative AIC. (B) At stage 5, stage 2 expression has a large negative AIC, while upstream gene distance is only slightly negative, indicating that the stage 2 expression is likely to be the most informative interaction.

**Fig S3:**
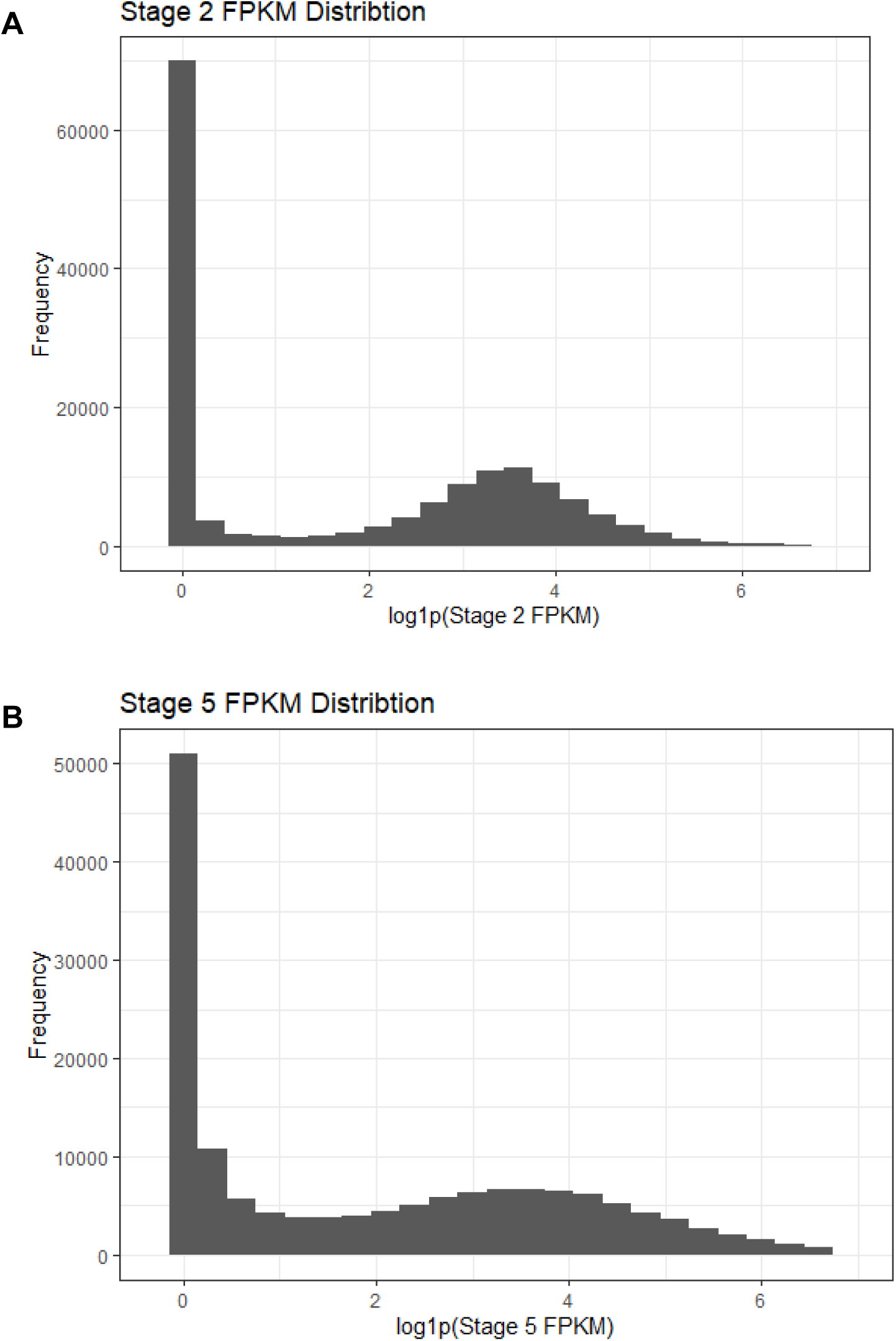
Transcript abundance distributions (in FPKM) are bimodal at both stages. We produced a histogram of the FPKM of all species at stage 2 (A) and stage 5 (B). At both stages, a peak appears at zero and approximately e∧3.5.

## References

Adkins, Nicholas L., Thomas A. Hagerman, and Philippe Georgel. 2006. “GAGA Protein: A Multi-Faceted Transcription Factor.” Biochemistry and Cell Biology = Biochimie et Biologie Cellulaire 84 (4): 559–67.

Alexa, Adrian, and Jorg Rahnenfuhrer. n.d. “Gene Set Enrichment Analysis with topGO.” https://bioconductor.org/packages/release/bioc/vignettes/topGO/inst/doc/topGO.pdf.

Arnosti, D. N., S. Barolo, M. Levine, and S. Small. 1996. “The Eve Stripe 2 Enhancer Employs Multiple Modes of Transcriptional Synergy.” Development 122 (1): 205–14.

Atallah, Joel, and Susan E. Lott. 2018. “Evolution of Maternal and Zygotic mRNA Complements in the Early Drosophila Embryo.” PLoS Genetics 14 (12): e1007838.

Bailey, Timothy L., Mikael Boden, Fabian A. Buske, Martin Frith, Charles E. Grant, Luca Clementi, Jingyuan Ren, Wilfred W. Li, and William S. Noble. 2009. “MEME SUITE: Tools for Motif Discovery and Searching.” Nucleic Acids Research 37 (Web Server issue): W202– 8.

Baroux, C., D. Autran, C. S. Gillmor, D. Grimanelli, and U. Grossniklaus. 2008. “The Maternal to Zygotic Transition in Animals and Plants.” Cold Spring Harbor Symposia on Quantitative Biology 73: 89–100.

Bashirullah, A., R. L. Cooperstock, and H. D. Lipshitz. 2001. “Spatial and Temporal Control of RNA Stability.” Proceedings of the National Academy of Sciences of the United States of America 98 (13): 7025–28.

Bownes, M. 1975. “A Photographic Study of Development in the Living Embryo of Drosophila Melanogaster.” Journal of Embryology and Experimental Morphology 33 (3): 789–801.

Briscoe, James, and Stephen Small. 2015. “Morphogen Rules: Design Principles of Gradient-Mediated Embryo Patterning.” Development 142 (23): 3996–4009.

Campos-Ortega, J. A., and V. Hartenstein. 1985. The Embryonic Development of Drosophila Melanogaster. Springer.

Cartwright, Emily L., and Susan E. Lott. 2019. “Evolved Differences in Cis and Trans Regulation between the Maternal and Zygotic mRNA Complements in the Drosophila Embryo.” Genetics 216 (3): 805–21.

Chenevert, M., Miller, B., Karkoutli, A., Rusnak, A., Lott, S., & Atallah, J. (2022). “The early embryonic transcriptome of a Hawaiian *Drosophila* picture-wing fly shows evidence of altered gene expression and novel gene evolution.” Journal of Experimental Zoology Part B: Molecular and Developmental Evolution, 338, 277–291. 10.1002/jez.b.23129

Chojnacki, Szymon, Andrew Cowley, Joon Lee, Anna Foix, and Rodrigo Lopez. 2017. “Programmatic Access to Bioinformatics Tools from EMBL-EBI Update: 2017.” Nucleic Acids Research 45 (W1): W550–53.

Cui, Jun, Caroline V. Sartain, Jeffrey A. Pleiss, and Mariana F. Wolfner. 2013. “Cytoplasmic Polyadenylation Is a Major mRNA Regulator during Oogenesis and Egg Activation in Drosophila.” Developmental Biology 383 (1): 121–31.

Eichhorn, Stephen W., Alexander O. Subtelny, Iva Kronja, Jamie C. Kwasnieski, Terry L. Orr-Weaver, and David P. Bartel. 2016. “mRNA poly(A)-Tail Changes Specified by Deadenylation Broadly Reshape Translation in Drosophila Oocytes and Early Embryos.” eLife 5 (*July*). 10.7554/eLife.16955.

Ellison, Christopher E., and Doris Bachtrog. 2013. “Dosage Compensation via Transposable Element Mediated Rewiring of a Regulatory Network.” Science 342 (6160): 846–50.

Gramates, L. Sian, Steven J. Marygold, Gilberto Dos Santos, Jose-Maria Urbano, Giulia Antonazzo, Beverley B. Matthews, Alix J. Rey, et al. 2017. “FlyBase at 25: Looking to the Future.” Nucleic Acids Research 45 (D1): D663–71.

Heinz, Sven, Christopher Benner, Nathanael Spann, Eric Bertolino, Yin C. Lin, Peter Laslo, Jason X. Cheng, Cornelis Murre, Harinder Singh, and Christopher K. Glass. 2010. “Simple Combinations of Lineage-Determining Transcription Factors Prime Cis-Regulatory Elements Required for Macrophage and B Cell Identities.” Molecular Cell 38 (4): 576–89.

Heyn, Patricia, Martin Kircher, Andreas Dahl, Janet Kelso, Pavel Tomancak, Alex T. Kalinka, and Karla M. Neugebauer. 2014. “The Earliest Transcribed Zygotic Genes Are Short, Newly Evolved, and Different across Species.” Cell Reports 6 (2): 285–92.

Hill, Tom, and Andrea J. Betancourt. 2018. “Extensive Exchange of Transposable Elements in the Drosophila Pseudoobscura Group.” Mobile DNA 9 (June): 20.

Jaeger, Johannes, Manu, and John Reinitz. 2012. “Drosophila Blastoderm Patterning.” Current Opinion in Genetics & Development 22 (6): 533–41.

Levine, Mike. 2010. “Transcriptional Enhancers in Animal Development and Evolution.” Current Biology: CB 20 (17): R754–63.

Li, Jian, and David S. Gilmour. 2013. “Distinct Mechanisms of Transcriptional Pausing Orchestrated by GAGA Factor and M1BP, a Novel Transcription Factor.” The EMBO Journal 32 (13): 1829–41.

Ludwig, Michael Z., Arnar Palsson, Elena Alekseeva, Casey M. Bergman, Janaki Nathan, and Martin Kreitman. 2005. “Functional Evolution of a Cis-Regulatory Module.” PLoS Biology 3(4): e93.

Ludwig, M. Z., N. H. Patel, and M. Kreitman. 1998. “Functional Analysis of Eve Stripe 2 Enhancer Evolution in Drosophila: Rules Governing Conservation and Change.” Development 125 (5): 949–58.

Medioni, Caroline, Kimberly Mowry, and Florence Besse. 2012. “Principles and Roles of mRNA Localization in Animal Development.” Development 139 (18): 3263–76.

Mishra, Krishnaveni, Vivek S. Chopra, Arumugam Srinivasan, and Rakesh K. Mishra. 2003. “Trl-GAGA Directly Interacts with Lola like and Both Are Part of the Repressive Complex of Polycomb Group of Genes.” Mechanisms of Development 120 (6): 681–89.

Obbard, Darren J., John Maclennan, Kang-Wook Kim, Andrew Rambaut, Patrick M. O’Grady, and Francis M. Jiggins. 2012. “Estimating Divergence Dates and Substitution Rates in the Drosophila Phylogeny.” Molecular Biology and Evolution 29 (11): 3459–73.

Omura, Charles S., and Susan E. Lott. 2020. “The Conserved Regulatory Basis of mRNA Contributions to the Early Drosophila Embryo Differs between the Maternal and Zygotic Genomes.” PLoS Genetics 16 (3): e1008645.

R Core Team. 2014. “R: A Language and Environment for Statistical Computing.” Vienna, Austria: R Foundation for Statistical Computing. http://www.R-project.org/.

Russo, C. A., N. Takezaki, and M. Nei. 1995. “Molecular Phylogeny and Divergence Times of Drosophilid Species.” Molecular Biology and Evolution 12 (3): 391–404.

Russo, Claudia A. M., Beatriz Mello, Annelise Frazão, and Carolina M. Voloch. 2013. “Phylogenetic Analysis and a Time Tree for a Large Drosophilid Data Set (Diptera: Drosophilidae): Drosophilid Timescale.” Zoological Journal of the Linnean Society 169 (4): 765–75.

Sallés, F. J., M. E. Lieberfarb, C. Wreden, J. P. Gergen, and S. Strickland. 1994. “Coordinate Initiation of Drosophila Development by Regulated Polyadenylation of Maternal Messenger RNAs.” Science 266 (5193): 1996–99.

Small, S., R. Kraut, T. Hoey, R. Warrior, and M. Levine. 1991. “Transcriptional Regulation of a Pair-Rule Stripe in Drosophila.” Genes & Development 5 (5): 827–39.

Tadros, Wael, Aaron L. Goldman, Tomas Babak, Fiona Menzies, Leah Vardy, Terry Orr-Weaver, Timothy R. Hughes, J. Timothy Westwood, Craig A. Smibert, and Howard D. Lipshitz. 2007. “SMAUG Is a Major Regulator of Maternal mRNA Destabilization in Drosophila and Its Translation Is Activated by the PAN GU Kinase.” Developmental Cell 12(1): 143–55.

Tadros, Wael, and Howard D. Lipshitz. 2009. “The Maternal-to-Zygotic Transition: A Play in Two Acts.” Development 136 (18): 3033–42.

Tsai, Shih-Ying, Yuh-Long Chang, Krishna B. S. Swamy, Ruei-Lin Chiang, and Der-Hwa Huang. 2016. “GAGA Factor, a Positive Regulator of Global Gene Expression, Modulates Transcriptional Pausing and Organization of Upstream Nucleosomes.” Epigenetics & Chromatin 9 (July): 32.

Vastenhouw, Nadine L., Wen Xi Cao, and Howard D. Lipshitz. 2019. “The Maternal-to-Zygotic Transition Revisited.” Development 146 (11). 10.1242/dev.161471.

Vazquez-Pianzola, Paula, Bogdan Schaller, Martino Colombo, Dirk Beuchle, Samuel Neuenschwander, Anne Marcil, Rémy Bruggmann, and Beat Suter. 2017. “The mRNA Transportome of the BicD/Egl Transport Machinery.” RNA Biology 14 (1): 73–89.

